# Recalcitrance of *Cannabis sativa* to *de novo* regeneration; a multi-genotype replication study

**DOI:** 10.1101/2020.06.23.167478

**Authors:** Adrian S. Monthony, Sean T. Kyne, Christopher M. Grainger, A. Maxwell P. Jones

## Abstract

*Cannabis sativa* is relatively recalcitrant to regeneration from somatic tissues, but several reports have been published demonstrating a response. Most reports show low levels of regeneration from somatic tissues, but a landmark publication by Lata *et al*. in 2010 reported regeneration from leaf explants with a 96% response rate, producing an average of 12.3 shoots per explant in a single, high-THC genotype. Despite the importance regeneration plays in plant biotechnology this protocol has not been used in subsequent papers in the decade since it was published, raising the concern that it is not reproducible. Many researchers are looking to build research programmes in this growing field, and it is important that the reproducibility and robustness of single-genotype *C. sativa* regeneration protocols undergo multi-lab validations to ensure they are reproducible across the species. Replication studies in this burgeoning field will help research groups avoid lost time and resources which arise from pursuing protocols that are not reproducible. Here we test the replicability of this protocol across 10 drug-type *C. sativa* genotypes. This protocol successfully induced callus in all 10 genotypes. Callus size and appearance substantially differed among cultivars, with the most responsive genotype producing 6-fold more callus than the least responsive genotype. However, the most successful shoot induction medium developed in the 2010 paper failed to induce regeneration in any of the cultivars tested, resulting in the eventual necrosis of the calli. Based on this replication study, it is evident that the existing regeneration protocol is not robust and could not be replicated in any of the 10 genotypes tested.

## 2 Introduction

Plant tissue culture provides the foundation for advanced biotechnological techniques such as protoplast systems, microspore culture for double haploid production, transgenics, genome editing, and other important tools which have yet to be explored in *Cannabis sativa* L. Due to prohibition, early micropropagation studies of Cannabis were limited in number and scope. As a result, the majority of past research relied on single drug-type genotypes or on less regulated industrial hemp (*Cannabis sativa* with <0. 0.03% w/w THC in the flowering heads) [1] cultivars as a proxy for drug type Cannabis. These early studies found that both drug-type Cannabis (high THC and/or high CBD) and industrial hemp can be maintained *in vitro* [2–4] and that *in vitro* grown *C. sativa* display comparable chemical and physical profiles to greenhouse-grown counterparts [2], with no significant effect on the cannabinoid contents of high-THC genotypes [4,5]. Regulations surrounding the production and consumption of Cannabis have since relaxed and the opportunity for biotechnological expansion in the Cannabis industry has increased. However, without robust and reproducible *in vitro* regeneration systems these technologies cannot be applied.

Regeneration systems are important to the application of plant biotechnologies. In Cannabis, the majority of micropropagation studies reporting *in vitro* regeneration in *C. sativa* rely on shoot multiplication from existing meristems found in the apical and axillary buds. Numerous protocols report regeneration from meristematic tissues at levels greater than 70% [4,6–8]. The most rapid *in vitro* multiplication methods of *C. sativa* are those which rely on existing meristems found in the nodal tissues of vegetative cuttings. *In vitro* cultures of these tissues can be used to establish multiple shoot cultures (MSC). The most successful protocols report between 9 and 13 shoot explants per nodal segment [4,7]. Unfortunately, most existing protocols were developed using a single high-THC genotype of *C. sativa*, and recent evidence suggests that methods developed using a single genotype are not replicable when tested on commercially available drug-type *C. sativa* genotypes and that multiplication rates are much lower, around 2.2 shoots per explant [9]. In addition to low regeneration rates, MSC is unsuitable for some biotechnological applications, and cannot compete with the relative ease of traditional vegetative propagation methods for large-scale commercial plant propagation.

Plant biotechnologies such as inter-specific hybridization via protoplast fusion or targeted genome editing through the use of CRISR-CAS9 (clustered regularly interspaced short palindromic repeats) have been well established in many species. However, these technologies have not been successfully applied in *C. sativa* due to a lack of reliable and robust regeneration protocols that are required to regenerate plants from somatic cells or protoplasts [10]. In addition to applications in biotechnologies, regeneration from somatic tissues would increase multiplication rates due to the wider range of responsive tissues compared to MSCs, which rely only on limited nodal tissues. While most published methods using non-meristematic tissue of Cannabis report low levels of regeneration [3,5,11–14], the use of a three-part regeneration system by Lata *et al*. [15] has shown success in leaf explants from a high-THC genotype, MX. In this protocol, the authors report optimized callogenesis on Murashige & Skoog (MS) media with 0.5 μM of α-naphthaleneacetic acid (NAA) and 1.0 μM thidiazuron (TDZ). Regeneration was most successfully induced by transferring cultures to MS media with 0.5 μM TDZ, resulting in 96.6 % of cultures responding with an average of 12.3 shoots per culture [15]. Rooting was subsequently achieved on half-strength MS and 2.5 μM indole-3-butyric acid (IBA) [15]. While only tested on MX, the authors state that “… the methodology described in this study may also be used for the propagation of male plants and any other genotypes of this [*C. sativa*] species (p. 1630)” [18]. However, despite the publication of this ostensibly successful protocol over a decade ago and the implications it has for advances in Cannabis biotechnology, it has not since been replicated in the literature by any independent research groups, including the original authors.

There has been an increasing awareness of what is dubbed the ‘reproducibility crisis’ in scientific literature and over 50% of biologists have reported that they have failed to reproduce published research [16]. This crisis has been attributed to structures in academia that base hiring of faculty, promotion and tenure on the novelty, impact and volume of published literature and journals who strive to publish “groundbreaking” research and who do not publish validation or replication studies, especially those studies showing negative results [16–18]. The incentivization of novel results arising from clean research narratives has been linked to incomplete reporting of methodologies and the omission of negative results [17] and with a rising number of retractions occurring each year [19] there is a case for more accountability through replication studies and ‘open data’ initiatives. Researchers often suffer from the prevailing narrative that volume of work is indicative of a high quality researcher [18]. A bellwether of this pressure to “publish or perish” are data obtained from the French CNRS (Centre national de la recherche scientifique) which showed that evolutionary biologists hired by the Centre in 2013 had published on average twice as many articles, that those hired in 2005 [20]. These aforementioned incentives promote novel results with succinct storylines at the expense of robust reporting of methodologies. The systemic challenges associated with publishing replication studies have eroded accountability and transparency in scientific literature. Publication of replication studies is necessary and can prove especially beneficial to new fields such as Cannabis micropropagation where the body of literature is limited and based on a narrow genetic base.

As the regulatory landscape evolves to facilitate research, there is growing interest in Cannabis regeneration and an increasing need for reliable methods (**Fig 1**). Despite the growing interest, the number of drug-type genotypes used in protocol development remains low, not exceeding 3 genotypes [5,14,21,22]. In stark contrast, regeneration studies on commercial hemp varieties can boast up to 12 genotypes [11]. Generalizing conclusions based on single-genotype studies have faced scrutiny as mounting evidence shows Cannabis responds in a genotype specific manner *in vitro*, yet few replication studies have been conducted [9,23,24]. The objective of this study was to replicate the Lata *et al*., 2010 methods across 10 different genotypes to assess the reproducibility of this protocol and the variability in genotypic response.

**Fig 1.**
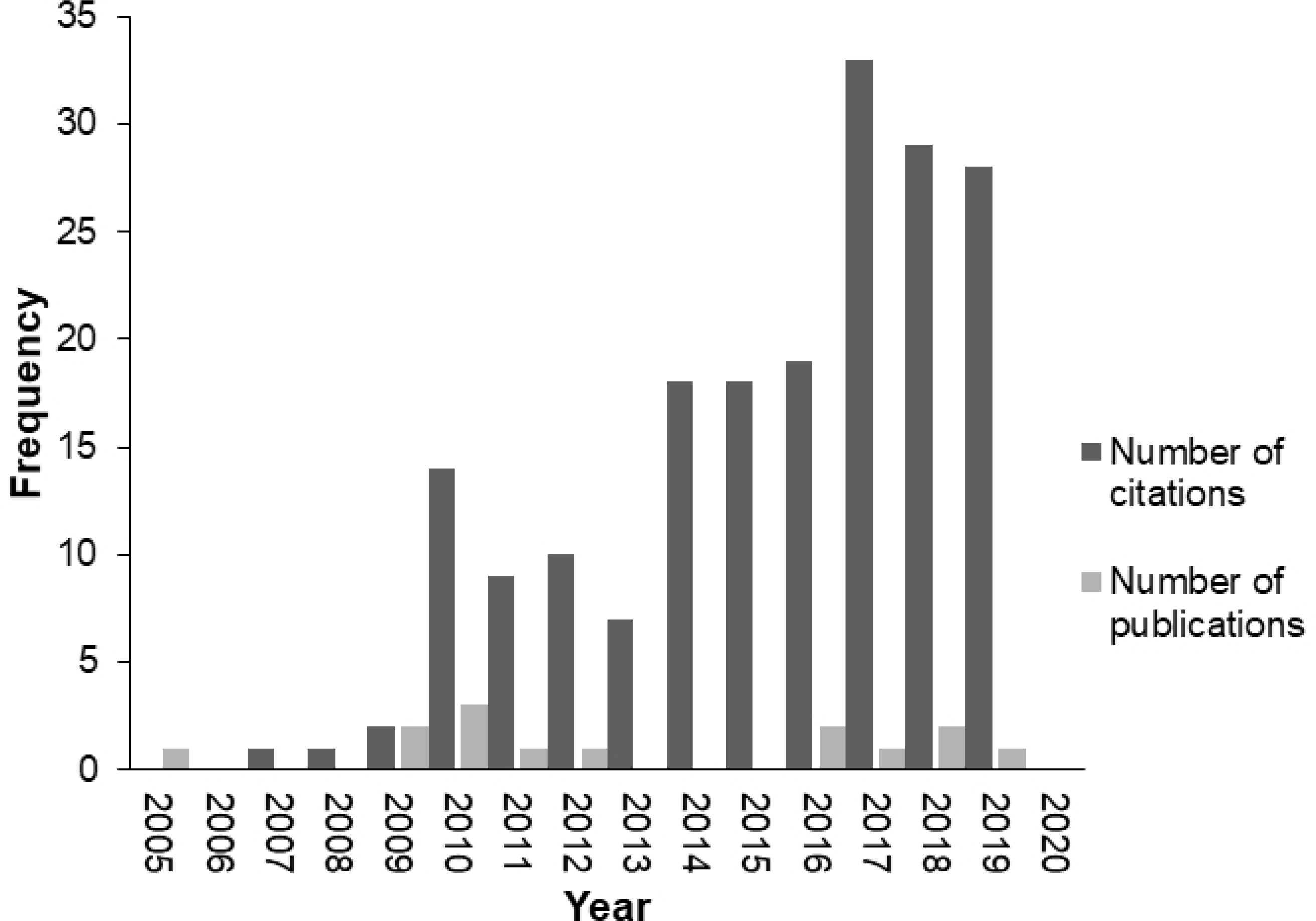
Annual citation frequency and number of new publications on regeneration in *Cannabis sativa*. The annual citation frequency on regeneration in *C. sativa* continues to grow, indicating a growing interest in the *in vitro* regeneration of Cannabis, however new publications on regeneration in Cannabis are not keeping pace. Search Topic: “Cannabis sativa” AND “regeneration” AND “in vitro” on the Web of Science database. Data obtained from Web of Science on March 17, 2020 using Microsoft Edge®.

## 3 Materials & Methods

### 3.1 Regeneration from Leaf-Derived Explants

#### 3.1.1 Plant Material

Leaf explants were excised from two *in vitro* grown drug-type genotypes. In the first experiment, two genotypes were used; GRC (clonal), and a heterogeneous seed-derived variety, RTG. After these two genotypes failed to respond as reported, the protocol was repeated using a more comprehensive genetic pool of 10 unique genotypes. For the second replication of the experiment, GRC and RTG were selected again, alongside 8 additional clonal genotypes, U22, U31, U37, U38, U42, U61, U82 and U91. All source material came from female explants cultured in We-V boxes (We Vitro Inc., Guelph, ON) each containing between 8 and 12 explants, which had spent a minimum of 6 months in *in vitro* culture. Cultures were maintained on semisolid medium composed of DKW nutrients (Phytotechnology Laboratories, KS, USA), 3 % sucrose and 6 g/L agar for 4 weeks in a controlled atmosphere growth chamber, under a 16-hr photoperiod at 25 °C. To ensure consistent sampling, leaves were selected based on their maturity, and ability to accommodate uniform 1 cm^2^ squares. As a result, young, fully emerged leaves found no lower than 3 nodes below the apical shoot were used.

#### 3.1.2 Media Preparation

The control medium (MS-0) consisted of MS (M524; Phytotechnology Laboratories, KS, USA) nutrients, 3% sucrose, 0.8% (w/v) type E agar (Sigma Aldrich, St. Louis, MO), and pH adjusted to 5.7 using 1 M NaOH and 1 M HCl. The callus induction medium (hereafter referred to as LT-C) consisted of MS (M524; Phytotechnology Laboratories) nutrients, 3% sucrose, 0.8% type E agar (w/v) (Sigma Aldrich), 0.5 µM NAA (Sigma Aldrich), and 1.0 µM TDZ (Caisson Laboratories, Inc., Smithfield, UT) adjusted to a pH of 5.7. The shoot proliferation medium (hereafter referred to as LT-S) consisted of MS (M524; Phytotechnology Laboratories) nutrients, 3% sucrose, 0.8% type E agar (w/v) (Sigma Aldrich), 0.5 µM TDZ (Caisson) and adjusted to a pH of 5.7. Each glass culture vessel (9.05 cm high x 5.8 cm diameter baby food jars; Phytotechnology Laboratories) contained 25 mL of media which were sterilized by autoclaving for 20 minutes at 121°C and 18 PSIG.

#### 3.1.3 Callogenesis

The leaf explants were excised into 1 cm^2^ squares using a sterile scalpel in a biological safety cabinet. All leaf explants were taken along the central axis of the leaflet, so as to include mid-rib tissues (**Fig 2A)**. Leaf explants did not include petioles. The abaxial side of one explant was carefully placed onto the medium in each glass jar culture vessels and gently tapped down to the surface of the media to maximize contact and mitigate unwanted movement while handling. Ten explants of each genotype were assigned to each treatment media (*n=10*). The capped jars were wrapped twice with Micropore™ tape (3M™, St. Paul, MN), randomized, and placed together in a controlled atmosphere growth chamber, under a 16-hr photoperiod at 25 °C. Photosynthetically active radiation (PAR) and light spectral data (**Fig S1**) were obtained using an Ocean Optics Flame Spectrometer (Ocean Optics, FL, USA). The average PAR of the experimental area was 41 ± 4 μmol s^-1^ m^-2^. Explants were maintained under these conditions for 8 weeks and checked weekly for callus development. Parameters measured included callus mass and percent response.

**Fig 2.**
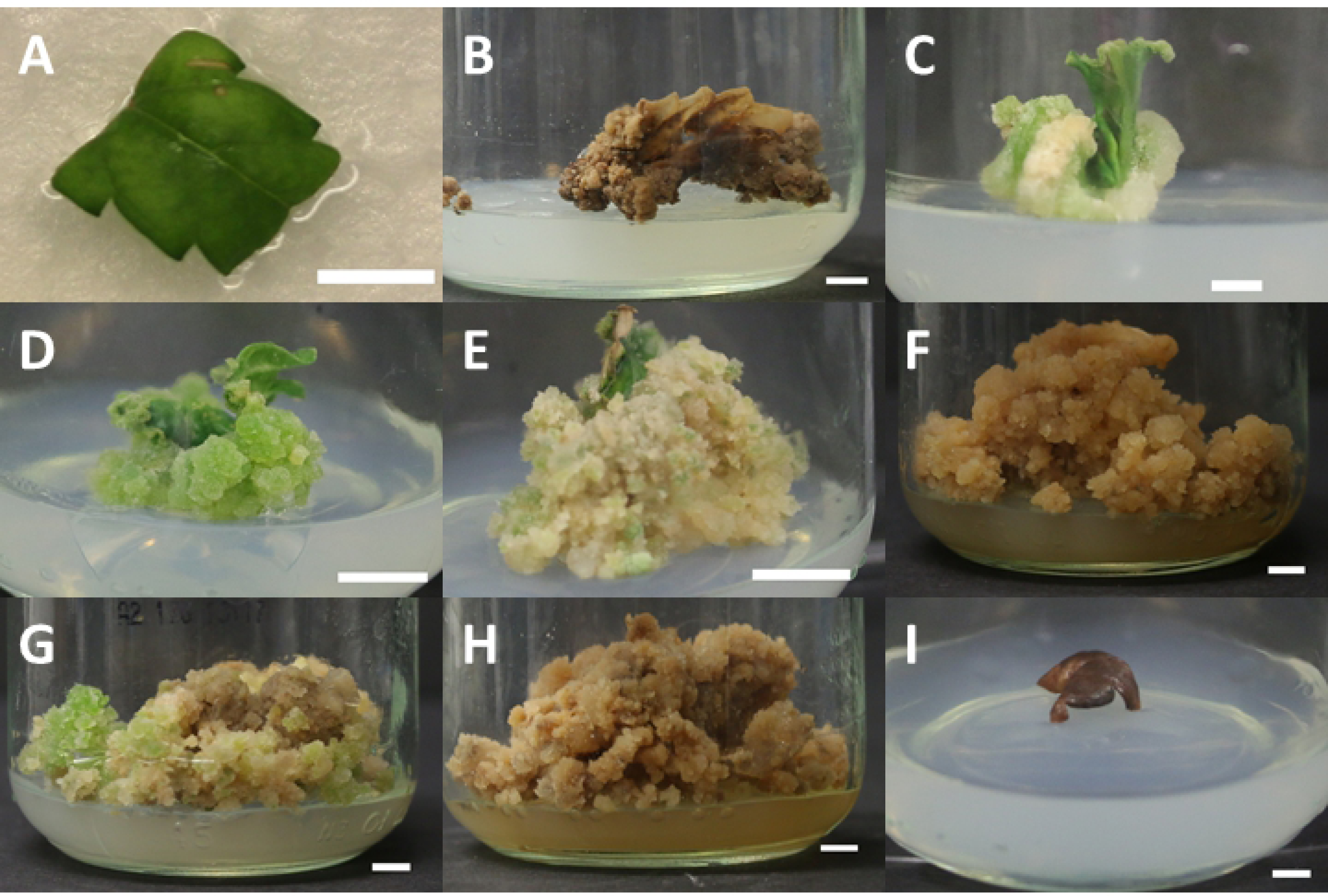
Explant and callus morphology over the course of callus and shoot induction phase of the replication study. A) Representative photograph of leaf explants of *C. sativa* cv. U82 used in the replication study, midribs but not petioles are included from the ∼ 1 cm^2^ leaflet. B) Necrotic callus of *C. sativa* cv. RTG after 4 months on LT-S medium. C) Cream coloured, friable callus of *C. sativa* cv. U91, one month after culture LT-C medium. D) Green, nodular callus of *C. sativa* cv. U61, one month after culture LT-C medium. E) Friable callus of *C. sativa* cv. RTG, two months after culture on LT-C medium, prior to transfer to shoot induction medium. F) Callus browning in *C. sativa* cv. U61 two months after transfer of callus onto LT-S medium. G) Some two month old callus on LT-S media showed only slight signs of browning with creamy and green nodular callus still present, pictured: cv. U82. H) Browning was ubiquitous across all cultivars following four months on LT-S medium, as seen by in cv. 82. I) Control leaf explants cultured on MS-0 media failed to callus, as shown by brown and dead leaf explant from cv. GRC. Scale bar in all photographs is 0.5 cm.

**Fig S1. Light spectrum used for *in vitro* growth of plant materials**.

A representative light spectrum of the lighting used in the controlled environment growth chamber. The average photosynthetically active radiation (PAR) over the experimental area was 41 ± 4 μmol s^-1^ m^-2^ using an OceanOptics. Average PAR was calculated using Excel ™.

#### 3.1.4 Shoot Initiation

For shoot initiation, 8-week-old callusing explants were moved from LT-C to LT-S medium and maintained under the previously described conditions. Callusing cultures were maintained on LT-S until shoots exceeded 2.5 cm in height, died (**Fig 2B)**, or for a duration of 4 months at which point they were considered non-responsive and were destroyed. Control explants from the callogenesis phase grown on MS-0 media which were still living were subcultured to baby food jars containing new MS-0 media following 8 weeks of culture. Parameters measured at this stage were the percent regeneration and number of explants produced per culture.

### 3.2 Simple Sequence Repeat (SSR) Analysis of Cultivars

#### 3.2.1 Plant Material

Fresh stem tissue obtained from in vitro grown explants was used for all DNA extractions. For each clonal genotype (U22, U31, U37, U38, U42, U61, U82, U91 and GRC) three, 1 cm stem segments each obtained from three separate clonal explants were pooled together. Due to the heterozygous nature of the RTG plants, a 2 cm segment was taken from each of the twelve *in vitro* explants used in both experiments and each of these 12 samples were processed separately. The approximate weight of each of these fresh tissue samples was 100 mg.

#### 3.2.2 DNA Extraction

DNA extraction was performed using NucleoSpin® Plant II Mini (Macherey-Nagel, Dürin, Germany) as per manufacturer’s instructions with minor modifications. The following modifications were made to the manufacturer’s procedure: PL1 and RNase A buffers were added to tissue samples in a 2.0 mL microcentrifuge tube prior to homogenizing. Samples were homogenized using a SPEX SamplePrep 1600MiniG® homogenizer (SPEX, Metuchen, NJ) at 1500 RPM for 1 minute. Following homogenization samples were briefly centrifuged to settle the lysate. Samples were gently resuspended and incubated for 15 minutes at 65 °C and extraction was subsequently performed as per manufacturer’s instructions. Extracted DNA was stored at −20 °C prior to SSR analysis.

#### 3.2.3 SSR Genotyping

Purified DNA was thawed at 4 °C and gently vortexed prior to quantification. DNA concentration and purity were measured with a NanoDrop ND-1000 spectrophotometer (Thermo Scientific, Waltham, MA). Eleven SSR markers **(Table 1)** developed by Alghanim & Almirall were used for genotyping [25]. PCR amplification was performed based on the Schuelke method [26]. Each PCR reaction consisted of: 3 µL 20% trehalose, 4.92 µL of molecular-grade H_2_O, 1.8 µL 10X PCR buffer (MgCl_2_), 3 mM dNTP mix, 0.12 µL of 4 μM M13-tailed forward primer, 0.48 µL of 4 μM reverse primer, 0.48 µL of 4 μM “universal” M13 primer labeled with VIC fluorescent dye (Applied Biosystems, Foster City, CA), 0.2 µL of 5.0 U µL^-1^ *taq* polymerase (Sigma Jumpstart™, Sigma-Aldrich, St. Louis, MO), and 3 L of template DNA for a total reaction volume of 15 µL. Amplification reactions were performed using thermocyclers (Eppendorf MasterCycler®, Hauppauge, NY). The conditions of the PCR amplification are as follows: 94 °C (5 min), then 30 cycles at 94 °C (30 s)/56 °C (45 s) /72 °C (45 s), followed by 8 cycles at 94 °C (30s)/53 °C (45S)/ 72 °C (45 s), followed by a final extension at 72 °C for 10 minutes. Following the PCR reaction, plates were held at 4 °C at the cycle’s end and subsequently stored at −20 °C until sequencing. Fragment analysis of the completed PCR products was done using an Applied Biosystems® 3500 Genetic Analyzer (ThermoFisher, Waltham, MA). A dye-labeled size standard (GeneScan 500-LIZ, Life Technologies, Burlington, ON, Canada) was used as the internal size standard, and PCR fragment sizes were determined using a DNA fragment analysis software (GeneMarker, SoftGenetics LLC, State College PA).

**Table 1.**
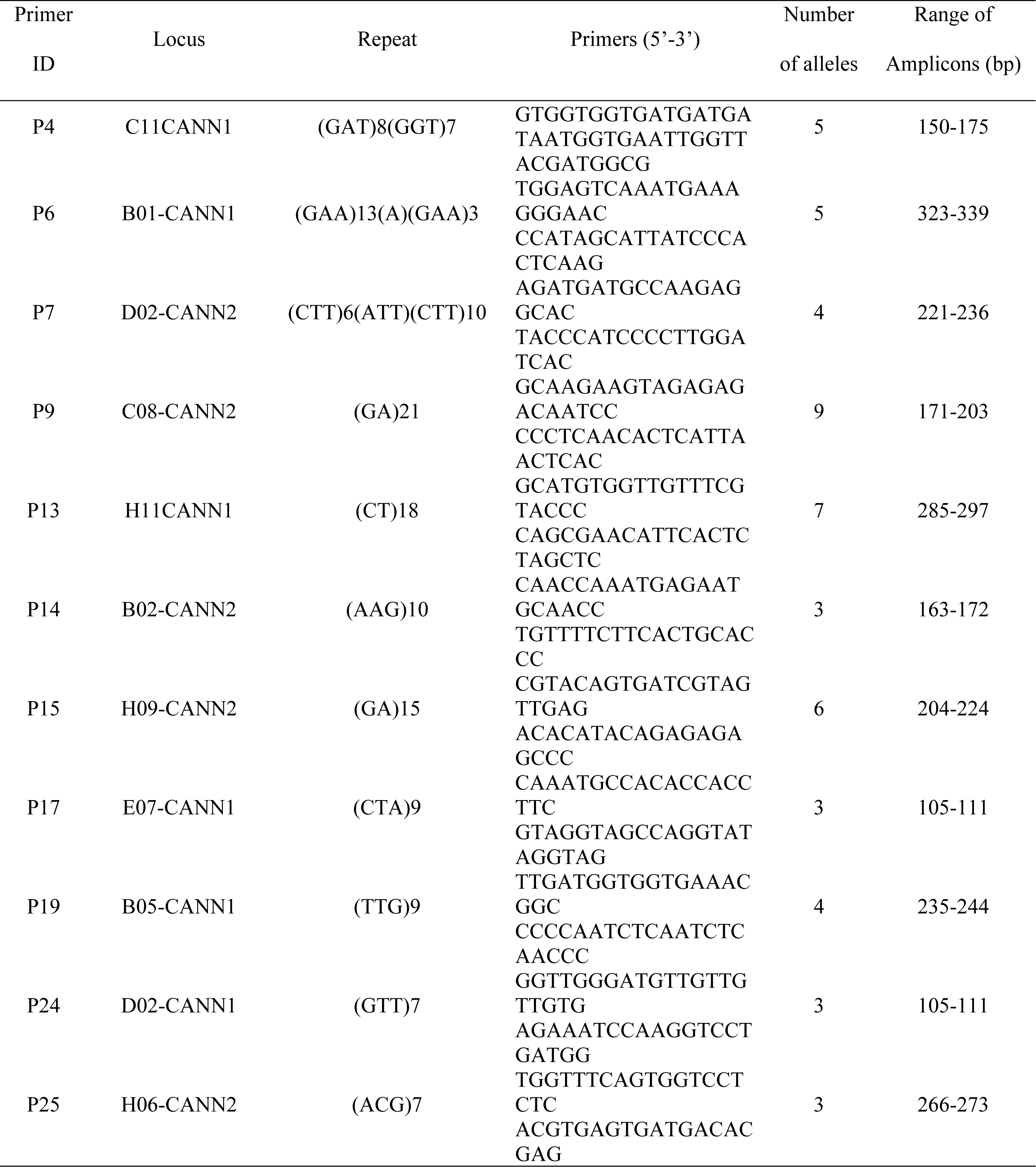
SSR loci and the primers used in this study, as first reported by Alghanim & Almirall [25]. Number of alleles and range of amplicon sizes obtained from experimental results.

#### 3.2.4 SSR Data Processing

SSR data (available on OSF database: https://osf.io/skd2f/?view_only=2bb948dec29a4a1db3ad4cf7eaece9bb) was analyzed and sorted using GeneMarker (SoftGenetics LLC, State College PA). Results of the SSR allele-call analysis were organized via Microsoft Excel (Microsoft Corp., WA, USA) and imported into GraphicalGenoTyping (GGT 2.0, Wageningen University, NL) and a relative genetic distance matrix was developed to compare pair-wise distance values (available OSF database: https://osf.io/skd2f/?view_only=2bb948dec29a4a1db3ad4cf7eaece9bb). This matrix was visually represented as a NJ dendrogram **(Fig. 3)**.

**Fig 3.**
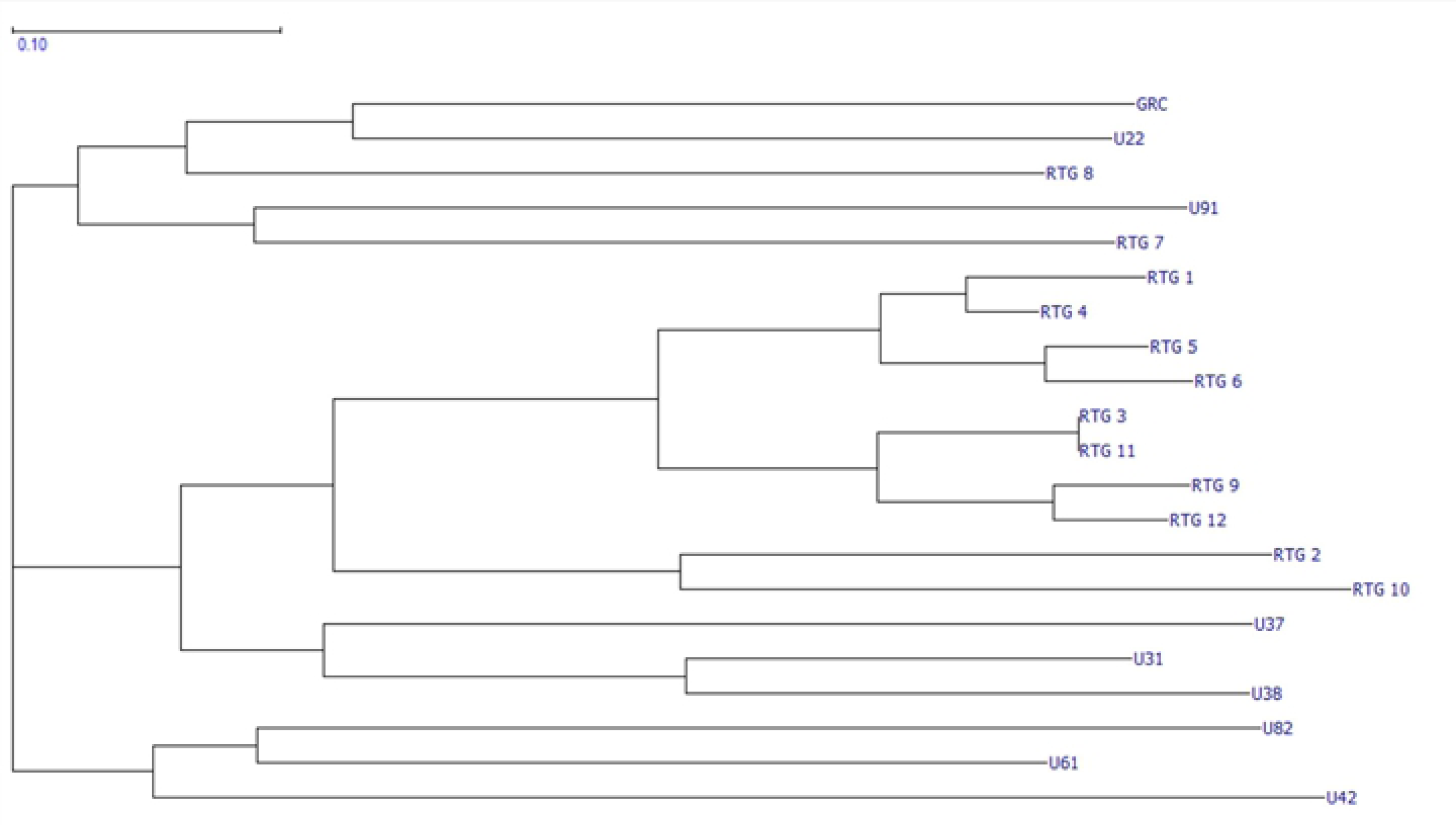
NJ dendrogram depicting relative genetic relatedness across selected cultivars. A visual representation of the relative genetic distance matrix presented as a NJ dendrogram. Data was analyzed using GeneMarker, organized in Excel, and visualized using GGT.

### 3.3 Experimental design and statistical analysis

Regeneration experiments were two-way cross-classified factorials with a completely random design testing two factors: genotype and media. The first experiment was a 2×2 factorial design testing two genotypes (GRC and RTG) against the previously described media MS-0 and LT-C according to the methods laid out by Lata *et al*. [22]. The experiment was replicated as a 10×2 two-way cross-classified factorial with the inclusion of 8 additional genotypes (U22, U31, U37, U38, U42, U61, U82, U91) in addition to GRC and RTG. Each treatment consisted of 10 experimental units (glass culture vessels; *n*=10), each containing one explant (sampling unit). All statistical analyses were performed using SAS Studio software (v9.4, SAS Institute Inc., Cary, NC, USA). The ANOVA was performed using PROC GLIMMIX and a means comparison of the callus mass of responding treatments was obtained using the LSMEANS statement (α=0.05). Missing data (due to contaminated cultures) were replaced with ‘.’ in the dataset (available on OSF Database: https://osf.io/94tae/?view_only=c484303807f540b0b94e31779b02253a) and were not processed by PROC GLIMMIX. Multiple comparisons were accounted for by a post-hoc Tukey-Kramer Test. were Visual presentation of the SAS data were prepared using Microsoft Excel® (Microsoft Corp., WA, USA).

## 4 Results & Discussion

### 4.1 Media proposed by Lata *et al*. does not induce regeneration in the 10 genotypes tested

The aim of this replication study was to assess the robustness of the high-frequency plant regeneration protocol from leaf derived callus put forward by Lata *et al*. in 2010 and assess the degree of genotypic variation in response. In our hands we found that the callus induction media proposed by Lata *et al*. [15] successfully induced callus in all 10 genotypes **(Fig 2C-E)**, however the subsequent shoot induction media failed to induce any regeneration in all 10 tested cultiars, resulting in browning of media, callus and death within four months of transfer to shoot proliferation medium LT-S (**Fig 2F-H)**. The callus induction rate was 100% across all 10 tested genotypes grown on the LT-C media (data available on OSF database: https://osf.io/kevu4/?view_only=1bfc16548f10495d9a20579111a076a0), which is comparable to the callusing rate reported by Lata *et al*. [15]. The earliest callus formation was observed within one week of induction, with all genotypes producing callus within two weeks. All genotypes continued to grow callus during the two-month callus induction phase **(Fig 2C-E)**. No callus formation was observed in any leaf explants in the 10 genotypes cultured on the control medium (MS-0). Within 2-month of initiation all control leaf explants on MS-0 turned brown and died **(Fig 2I)**. Callusing in the absence of PGRs is uncommon in Cannabis and Lata *et al*. similarly reported no callusing on MS salts without PGRs [15].

Callus formation and appearances have been previously been shown to vary across *C. sativa* genotypes. Early work from Mandolino & Ranalli [11] looking at the induction of embryogenic callus found that callusing was easily achieved in leaf tissues of 12 hemp genotypes. However, leaf tissues failed to regenerate in most cases, with only hypocotyls from 1 of the 12 tested genotypes regenerating [11]. Ślusarkiewicz-Jarzina *et al*. [3] similarly reported that callogenesis frequency and appearance varied in their screening of explant tissues for organogenic potential in 5 hemp genotypes. They reported that young leaves were most responsive to callusing with 52% of explants responding, however only 1.35% of the calli regenerated [3]. The results of our study confirm that the tested LT-C medium was more effective than previous protocols at inducing callogenesis, reaching 100% response across all 10 tested genotypes. This high response rate compared with previous callogenesis studies on hemp could indicate that drug-type genotypes are more responsive to callusing than some industrial hemp genotypes, or that the medium itself is more effective, and future studies should be undertaken to understand how industrial hemp and drug-type genotypes vary in their callogenesis response.

To date, few studies have been conducted on drug-type genotypes to observe the effect of genotype on callogenesis or regeneration. Piunno *et al*. [5] tested regeneration potential of three drug-type genotype and achieved callusing and low levels of regeneration in 2/3 of the genotypes, but it is unclear from this report if this was *de novo* regeneration or proliferation of existing meristems [5]. To date, this experiment represents the largest replication of an existing *C. sativa* regeneration protocol. While we found that callusing was obtained on LT-C medium across all 10 tested genotypes, rates of callogenesis were genotype specific (**Fig 4 & Fig S2)**. The average mass of callus differed significantly among genotypes with the most responsive, UP-601 producing 6-fold more callus than the least responsive genotype, RTG (6448 vs. 1180 grams; **Fig 4**). Callus appearance also varied across genotypes from a friable cream colour (**Fig 2C)** to hard nodular green callus (**Fig 2D**). These differences were apparent within a month of callus induction. Callus growth persisted for the duration of the callus induction phase (2 months; **Fig 2E)**. Our findings that callus formation is not uniform across the genotypes tested suggest that the response is not universal across *C. sativa* genotypes, which could also reflect regenerative capacity. A similar genotype effect was demonstrated by Wielgus *et al*.(2008) in industrial hemp who showed that while callogenesis response existed across the species, the ability to regenerate was genotype specific [13]. Similarly, Chaohua *et al*. found that the regeneration response was partly genotype dependent [27].

**Fig 4.**
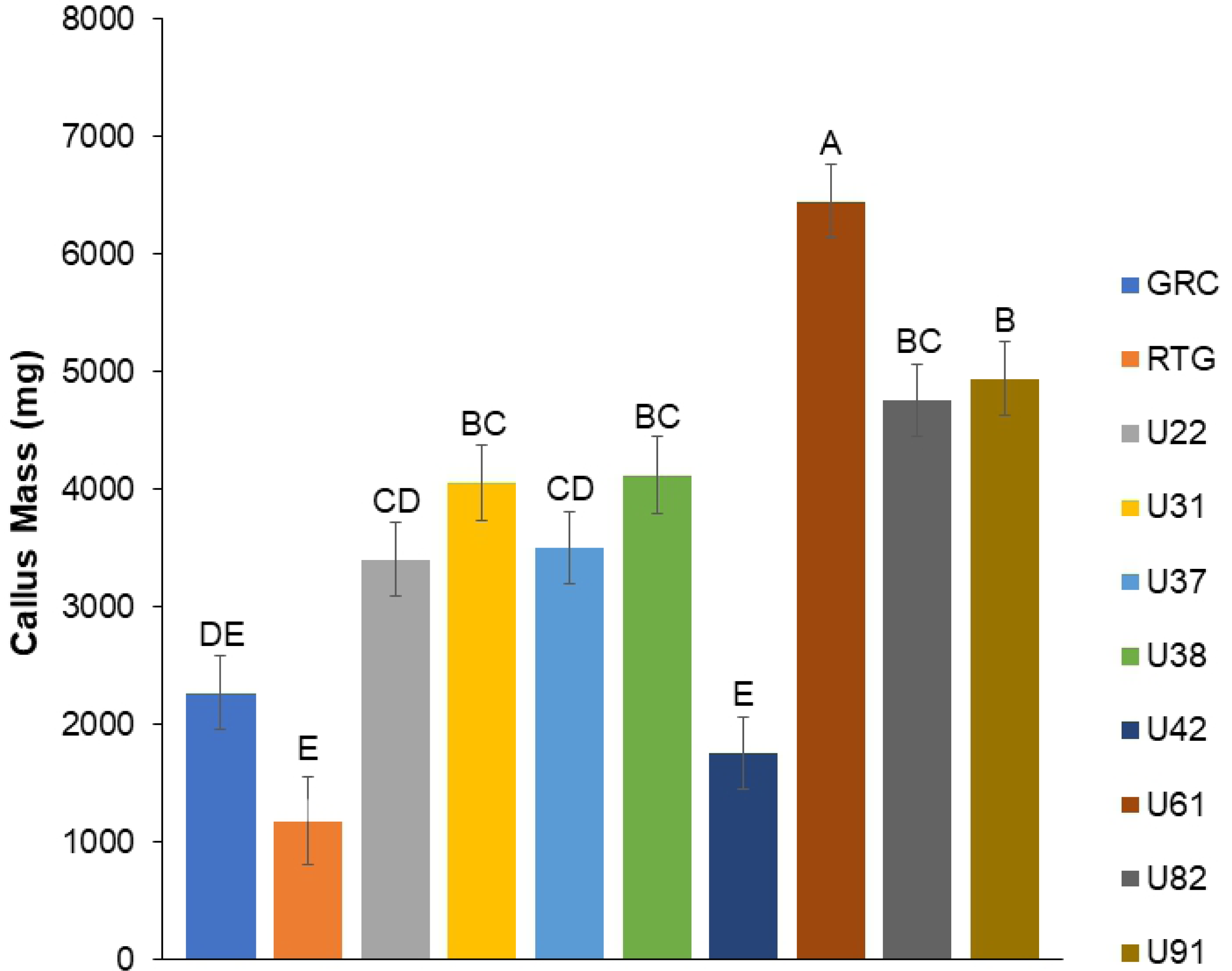
Average callus mass (mg) produced by a 1 cm by 1 cm leaf square following 2 months on LT-C media. All 10 cultivars showed a callus response. Control media contained no PGRs and did not induced callus in any of the tested cultivars. Same letters indicate that means were not significantly different at p=0.05 as determined by a Tukey-Kramer multiple comparisons test.

**Fig S2. Average callus mass (mg) produced by a 1 cm by 1 cm leaf square following 2 months on LT-C media in genotypes GRC and RTG**.

Callogenesis was first tested in two commercially available genotypes prior to a subsequent screening of the complete 10 cultivars. Callogenesis was achieved on MS media supplemented with 1.0 μM TDZ and 0.5 μM NAA. Control media contained no PGRs and did not induced callus in any of the tested. Same letters indicate that means were not significantly different at p=0.05 as determined by a Tukey-Kramer multiple comparisons test.

High callusing levels are not uncommon in Cannabis despite low to no regeneration from somatic callus. It has been suggested that these high callusing levels are due to high levels of endogenous auxins in the species, further supported by the strong apical dominance and rooting capacity of the species [7]. Regeneration of explants from somatic tissue requires the right auxin: cytokinin ratio. Auxin inhibitors have been shown to be more effective at reducing the auxin: cytokinin ratio rather than cytokinin supplementation as was done in this replication of *Lata et al*.’s method [15]. High endogenous levels of auxin may have played a role in why callusing was easily achieved across the 10 tested genotypes, but no regeneration was achieved. The variability in cultivars response to MS media with TDZ for callogenesis and regeneration was recently demonstrated by Chaohua *et al*. [27] whose callogenesis and shoot induction protocol used MS supplemented with TDZ and NAA for both callogenesis and shoot induction, rather than separate callogenesis and shoot proliferation media as proposed by Lata *et al*. [15]. Chahua *et al*. reported this media reduced the shoot induction period reported by Lata *et al*. from 2 months to 4 weeks, however rates of regeneration were considerably lower [27]. Smýkalová *et al*. [7] and Wróbel *et al*. [8] have also suggests that frequent subculture is required in order to maintain the regenerative potential of callus cultures of Cannabis, which was not done in the author’s original work and therefore not performed in this replication study. Work by Page *et al*. [9] has shown that the use of DKW media [28] may promote increased levels of callogenesis in some Cannabis genotypes, suggesting that media composition and genotype may interact to affect the formation of callus and subsequent regeneration potential.

Using the shoot proliferation medium proposed by Lata *et al*. [15] we failed to obtain regeneration. Within 2 months of culture onto the LT-S medium calli began producing phenolic compounds in the media **(Fig 2F)** indicative of stress, however some showed a pleateau in growth and no signs of stress (**Fig 2G)**. By 4 months on the LT-S media, most calli showed signs of necrosis and phenolic production (**Fig 2H)** while those not producing phenolics were were considered to be non-responsive to the shoot induction treatment as replicated from Lata *et al*. [15] and were destroyed. While genotypic variation is one explanation for our failure to obtain regeneration in the 10 tested genotypes, it could also be due to other factors that were not replicated exactly. One potential cause could be the use of *in vitro* grown leaves rather than greenhouse derived tissues as used by Lata et al., 2010. While *in vitro* leaves are generally more responsive to regeneration protocols in other species [29,30], it should be noted that physiological differences or the impact of surface disinfection used in the original protocol could account for the discrepancy of our results from those in the 2010 protocol. Additionally, there were a number of aspects of the protocol that were not well described in the original protocol, making the precision of some aspects of this replication study unknown, for example details around leaf explant selections and preparation (inclusion of midrib, petiole, leaf age) were scant, but play an important role in assuring the protocol can be accurately repoduced. As such, it is possible that the lack of reproducibility found here is a result of minor differences in our methodology however, irrespective of these differences we highlight that the protocol is difficult to reproduce.

### 4.2 SSR analysis indicates that each of the selected *C. Sativa* genotypes are genetically distinct

As shown in **(Fig. 3)**, the results of the SSR analysis identify all 10 cultivars as genetically unique. Cultivars are organized into three predominant clusters. As expected, the most closely related samples were found within the seed-derived lineage of RTG-X, however samples RTG-7 and RTG-8 present as outliers and lend support to the argument that Cannabis seed is highly heterogeneous. Clustering patterns also suggest genetic relatedness between U3X and RTG-X groups. The information displayed in **(Fig. 3)** demonstrates that the material used in this study represents genetically distinct genotypes of *C. Sativa*.

## 5 Conclusion

Since publication in 2010, there have been no independent research groups who have published a replication of the landmark study by Lata *et al*. on the regeneration of explants from leaf-derived callus of a single Cannabis genotype. This replication study is the first of its kind in the field of Cannabis micropropagation, while also expanding the scope of the original work to determine its replicability across 10 drug-type Cannabis genotypes. Our study shows that the most successful callus induction media (MS with 1.0 μM TDZ+0.5 μM NAA) proposed by Lata *et al*. induces callus growth in all 10 tested drug-type Cannabis genotypes, however the quantity of callus induction was found to be species-specific. These findings further contribute to growing evidence that micropropagation of Cannabis is highly genotype specific. We were unable to successfully induce regeneration by transferring callus to MS media supplemented with 0.5 μM TDZ in any of the 10 tested genotypes. The failure of this part of the method, which originally reported regeneration levels exceeding 96%, raises doubts about the original authors’ claims that this protocol can be used on any genotype of the species.

## Data Availability Statement

All data are available from the OSF database with the following DOI: 10.17605/OSF.IO/QVUG2

## 6 Acknowledgments

The authors gratefully acknowledge our industry partner, Hexo Corp. for the use of their plant material. The financial support of the Natural Sciences and Engineering Research Council of Canada (Grant No. RGPIN-2016-06252; awarded to AMPJ) is also gratefully acknowledged. Hexo Corp. (https://www.hexocorp.com/) and the Natural Sciences and Engineering Research Council of Canada (https://www.nserc-crsng.gc.ca/index_eng.asp) were not involved in study design, data collection and analysis, the decision to publish or the preparation of the manuscript.

